# Sequential critical periods support efficient local representation learning in a model of visual processing

**DOI:** 10.1101/2024.12.20.629674

**Authors:** Ariane Delrocq, Wu S. Zihan, Guillaume Bellec, Wulfram Gerstner

## Abstract

The emergence of abstract object representations in the mammalian ventral visual stream remains a central challenge for biologically plausible learning theories. While deep artificial networks trained via backpropagation achieve high performance, the algorithm lacks biological realism and conflicts with the staggered timeline of critical periods observed in cortical development. Here, we present a hierarchical model of the visual stream that learns invariant representations using only local synaptic plasticity rules modulated by lateral predictive signals. We demonstrate that imposing staggered critical periods — where plasticity windows open and close sequentially from V1 to inferotemporal cortex — significantly enhances representation quality for local learning rules, whereas it degrades performance in backpropagation-based networks. Furthermore, this sequential regime improves the learning efficiency in terms of number of synaptic updates, suggesting a metabolic advantage consistent with evolutionary constraints. We validate the functional utility of these acquired representations through reinforcement learning agents that successfully solve navigation and visual discrimination tasks without further fine-tuning of the visual encoder. These findings suggest that staggered critical periods are not merely a developmental constraint but a functional mechanism that enables efficient, local, and metabolically economical learning in hierarchical neural systems.

## Introduction

While visual input consists of high-dimensional retinal ‘pixel’ patterns, humans perceive the world as organized into objects like trees or coffee mugs. There are good reasons for the brain to transform sensory input patterns into more abstract representations. For example, different behavioral tasks might be learned more rapidly if the brain can rely on a common representation of the world. Along the ventral visual stream from early visual cortex V1 to higher visual areas in inferotemporal cortex (IT) the level of abstraction increases from simple edge-detectors in V1 [1] to face-sensitive and object-sensitive neurons in IT [2, 3, 4]. Cortical representations are not genetically prewired but adapt during development [5, 6] and can be altered by changes in input statistics [7, 8, 9, 10, 11, 12, 13].

The rules of synaptic plasticity that enable abstract representations to emerge across several steps of the cortical processing stream remain difficult to grasp [4]. Classic models of Hebbian plasticity [14], where the change of a synapse depends only on the state of pre- (pre) and postsynaptic (post) neurons (and potentially on the momentary state of the synapse itself) have been useful to describe the development of receptive fields (RFs) in models of V1 [15, 16, 17, 18, 19, 20, 21, 22, 23], but are not sufficient to explain the increase of abstraction found across the ventral visual stream [4]. Currently, the most powerful models of object representation in the mammalian ventral stream are multilayer neural networks optimized by gradient descent with the backpropagation algorithm [24]-[30]. However, the backpropagation algorithm is biologically not plausible [31, 32], even though variants of the algorithm may increase some aspects of bio-plausibility [33]-[45]. A second path proposed to reconcile learning in deep networks with biology is the concept of predictive coding [46]-[52]. Despite the partial success of a recent study [53], it has overall been difficult to arrive, with Hebbian or predictive learning rules, at a hierarchical representation defined such that receptive fields in deeper cortical areas are both more complex *and* more useful than in the first areas. Indeed, the performance of networks with biologically plausible Hebbian rules lags far behind that of deep artificial neural networks trained with backpropagation [29].

However, recent plasticity experiments have reported observations that cannot be explained by standard Hebbian learning rules. In particular, synaptic plasticity at basal dendrites of pyramidal neurons depends on input arriving in the apical dendrite [54, 55], potentially conveying predictive signals from other neurons located in a different information processing pathway [54] or a higher cortical area [55]. Here we use such predictive, plasticity-modulating apical input to induce representation learning in an Artificial Neural Network (ANN) model of the ventral visual stream. Each layer of the ANN represents an area in the visual processing stream. In addition to the usual feed-forward connections, our network also has lateral connections within each area. From an abstract perspective, the lateral predictive input materializes the idea that, within the *same* visual image, obstructed parts should be predictable from their surround, consistent with experimental results [49, 56, 57] - but predictions should not be possible across separate images. In a broader sense, our approach is linked to contrastive predictive coding (CPC) in machine learning [58] -[65] but avoids the known problems of backpropagation [31, 32].

With the backpropagation algorithm, synapses across the whole brain are optimized in parallel. This simultaneous change of all synapses is in contradiction with the presence of critical periods of plasticity [66], i.e., limited time windows during which a given cortical area is highly plastic and particularly sensitive to specific aspects of sensory stimuli [67, 68]. Windows of critical-period plasticity have been found to be staggered across different areas, potentially to first build the elements necessary for a later representation of relevant, but highly abstract features [69]. Language acquisition from auditory signals across multiple brain areas and time scales is a prominent example of this concept [70]. The opening and closing of critical periods involves multiple genetic, molecular, and cellular factors [69, 71] that enable the formation of new as well as changes of existing synapses. Inspired by the timeline observed in critical periods, here we use a model of learning in the ventral visual stream to study how representations can be learned sequentially over the hierarchy of areas from V1 to IT.

## 1 Modeling learning in the ventral visual stream

We describe the visual stream from V1 to V2, V4, posterior, central, and inferior temporal cortex by a convolutional neural network with six layers where each network layer corresponds to one cortical area [27, 53, 72]. The size of neuronal receptive fields (RFs) increases across areas. Neuron *i* in one of the areas has a firing rate

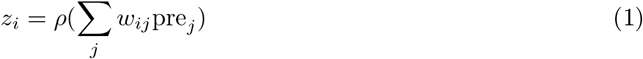

where pre_*j*_ is the rate of a neuron *j* in the preceding area and *w*_*ij*_ is the feedforward weight that connects a given area with the following area in the processing stream; the neuron’s frequency-current (f-I) curve is approximated by a rectified linear function *ρ*. Compared to the cortex, the architecture of our model is simplified, since we neglect backward connections and the detailed microcircuits of excitatory and inhibitory neurons within each area. However, we take into account that, within each area, feedforward input arriving mainly at the basal dendrites of pyramidal neurons has a different role than feedback signals arriving mainly in the apical dendrites [73], either directly or indirectly via higher-order thalamus [32, 73, 74]. For the sake of simplification, we consider in the model that the major source of feedback stems from neurons in the same cortical area (Fig. 1), but we also considered model variants where feedback arrives from higher areas.

**Figure 1.**
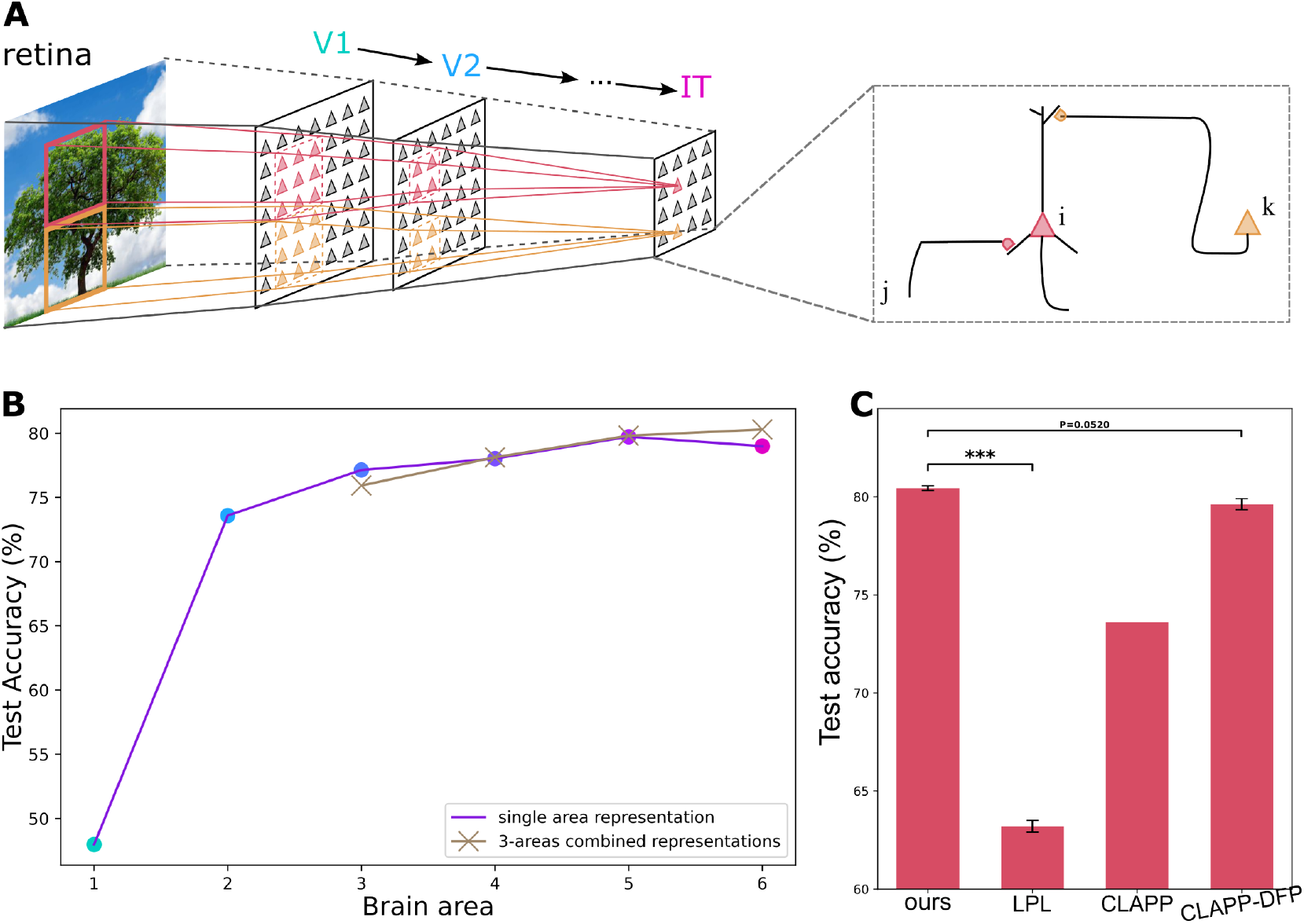
Model of synaptic plasticity and learning. **A.** Layered convolutional neural network representing areas V1, V2, V4, posterior, central, and inferior temporal (IT) cortex. Within each area, the network has lateral connections. If the image of a tree is projected onto the retina, a neuron (red triangle) with an RF in the red-framed square receives lateral input from other neurons (orange triangles) with non-overlapping RFs (orange square). Zoom: The plasticity of the feedforward connection *w*_*ij*_ from neuron *j* in the preceding area to neuron *i* (red) in IT depends on lateral input received by neuron *i* from other IT neurons *k* (orange). **B**. Performance on STL-10 (test accuracy using linear decoding of the representations) of our model with critical periods (CP) using representations from different brain regions. The usefulness of representations increases along the visual stream. Best accuracy is obtained by reading out from three IT areas in parallel (brown: readout from areas 1-3, 2-4, 3-5, 4-6). Since brain areas that read out the visual stream have access to multiple areas in IT[78], this is our reference model in the following. **C**. Performance on STL-10 (test accuracy using linear decoding of the representations) of our model with critical periods (corresponding to rightmost brown point in **B**.) compared to published models of bio-plausible learning: LPL [53], CLAPP [72], CLAPP with direct feedback [79]. The values of other models are taken from the original publications.

We model plasticity using the CLAPP learning rule [72], inspired by classic results of synaptic plasticity: the changes of a feedforward synapse from a presynaptic neuron *j* onto the basal dendrite of a postsynaptic neuron *i* are described by a Hebbian mechanism that depends on the firing rate pre_*j*_ of the presynaptic neuron as well as the state of the postsynaptic neuron [75, 76, 77]. In our model, the state of the postsynaptic neuron is characterized by two components. First, it depends on the local membrane potential post_*i*_ of the postsynaptic neuron which is a function post_*i*_ = *f*(*a*_*i*_) of the total *feedforward* input *a*_*i*_ = ∑_*j*_ *w*_*ij*_pre_*j*_ into the basal dendrite; second, it also depends on the total *lateral* input 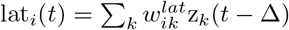 into the apical dendrite caused by the activity *z*_*k*_ of other neurons in the same area, where Δ denotes a small delay of reception and integration of the apical dendritic input. The change of the feedforward weights to basal dendrites is

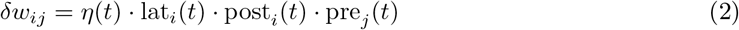

where *η*(*t*) denotes a learning rate. The lateral connections 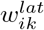 change with a Hebbian learning rule as in [72]. The direction of change (potentiation *δw*_*ij*_ > 0 or depression *δw*_*ij*_ < 0, Eq. 2) of a model synapse *w*_*ij*_ onto the *basal* dendrite depends on the *apical* input of the postsynaptic neuron, consistent with experiments [55]. Since we do not distinguish between excitatory and inhibitory neurons, all connections take positive or negative signs.

To illustrate the functionality of the learning rule, we focus on one of the areas in the processing stream and select two regions in that area (Fig. 1A). The Hebbian synaptic plasticity rule in Eq. (2) adjusts connectivity so that the lateral feedback input that arrives from region 2 onto an arbitrary neuron *i* in region 1 matches the feedforward input into neuron *i*. Indeed, during the stimulation with a static image the learning rule minimizes in each brain area the loss [72]

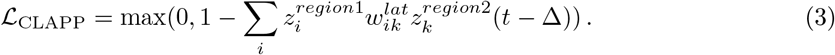

Thus, after learning, feedback input plays a predictive role, consistent with experiments [49, 56, 57]. The learning rate *η*(*t*) in Eq. (2) vanishes, if the match between feedforward and feedback input is good, but is large if the match is not good (Methods). The learning rate can therefore be considered as modulated by surprise [80, 81], i.e., learning is switched on if the expectation conveyed by lateral input does *not* match the actual input conveyed by the feedforward pathway. Such a switching process could be implemented by the coupling and decoupling of apical from basal dendrites at appropriate moments potentially controlled by thalamic input [82]. Switching the sign of plasticity during transitions between different images (Methods) avoids representational collapse. We conjecture that transitions between different objects in a natural scene are caused by large saccadic eye movements and trigger brain signals [83, 84] that relate views to each other [10, 85]. Alternative normalization schemes that avoid collapse are also possible [53, 86]. In summary, the generalized plasticity rule of Eq. (2) is, in a broad sense, in qualitative agreement with synaptic plasticity observed in cortical pyramidal neurons.

Our approach radically departs from previous work [45, 53, 55, 72] in two aspects. First, departing from [72], predictive input is generated, within each cortical area, by localized regions of cortical neurons at relatively long distances across the retinotopic map, rather than by the same neurons across a sequence of within-object micro-saccades with a tiny field of view. This conceptual shift enables us to better exploit the spatial structure of the visual input (Fig. 1A). Second, and more importantly, we model critical periods (CP) [66, 67, 68, 69] by implementing sequential learning, one area after the other, along the visual processing stream, as explained in the next section. We tested our learning rule, Eq. (2) with CPs, while stimulating the retina with thousands of images from the STL10 data base [87]. During training we neither provide the identity of objects nor do we allow any backpropagation of information from IT to lower areas. Nevertheless, the usefulness of the representation, as measured by linear decodability of the object identity, increases from model V1 to model IT (Fig. 1B), consistent with experimental findings that the level of abstraction is higher, and closer to object identity, in area IT [4] than in V1. Learning with CPs surpasses (Fig. 1C) earlier neuro-inspired modeling work at the interface to machine learning [72, 53]

## 2 Critical periods are compatible with local learning, but not with BackProp

We compared CP learning using the local area-specific loss function of Eq. (3) with end-to-end learning in a network of the same architecture. To enable a fair comparison, we used the same loss function, but applied only to area IT while the connections of all other areas are adjusted with the standard backpropagation algorithm (‘end-to-end training’). As expected, test accuracy using linear readout from the neural representations is higher for backpropagation than for the local synaptic rule (Figure 2A). However, when we combine the backpropagation algorithm with sequential CPs, the network performance gets worse, whereas the performance of the local synaptic plasticity rule improves (Fig. 2A). The reason is that, with end-to-end learning, the representations of all brain areas co-evolve so as to optimize the loss of the final area (IT); decoupling brain regions breaks the consistency between brain areas: during the V1 critical period, V1 learns to optimize its representations *for the higher areas that still have connections at their random initialization values*. After the end of this CP, V1 connections remain fixed, despite the fact that higher areas start learning and sending relevant feedback. In contrast, our local learning does not use any information from higher areas. Therefore, each region learns *independently of all higher areas*. To the contrary of the backpropagation case, it is beneficial to start learning higher layers only once the earlier layers are fixed, so that the former can build on consistent, rather than evolving, input from the latter. The fact that brain areas follow staggered CPs [69] indicates that the brain is unlikely to use (an approximation of) backpropagation algorithms during development.

**Figure 2.**
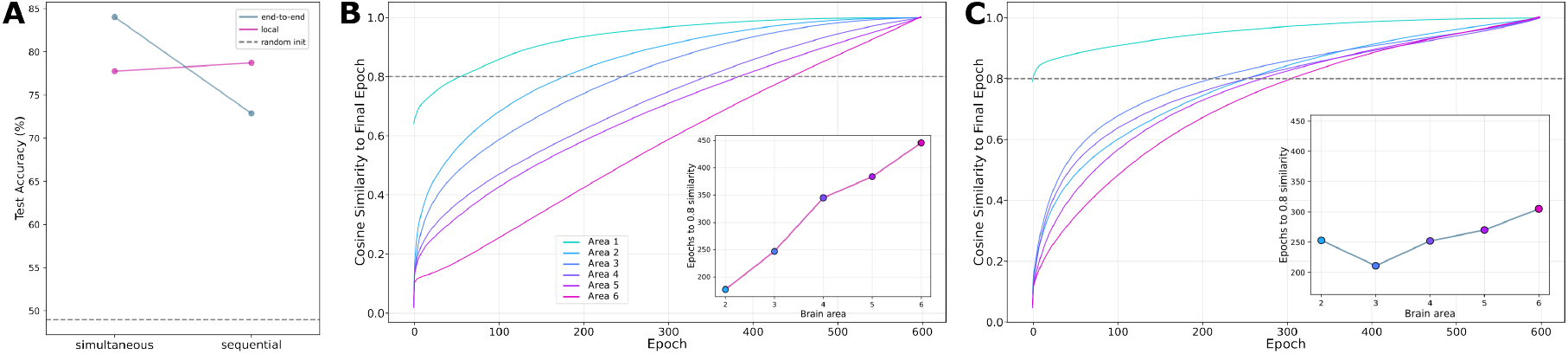
Sequential critical periods are compatible with local learning rules, but not with BackProp. **A.** We impose sequential critical periods by enabling plasticity in only one brain area at a time, with areas being plastic sequentially across the hierarchy. BackProp training suffers from imposed sequential learning while local learning benefits from it. Test accuracy from the neural representations (in last area for end-to-end, combined in last three areas for local, *cf* Methods), for the local and end-to-end models, in the standard parallel training case and in the case of imposed sequential training (300 epochs per area). **B**. and **C**. Learning naturally staggers sequentially across the hierarchy in layerwise (LW) but not BackProp learning, even with standard simultaneous training. Cosine similarity along training between each area’s synaptic weights and their respective final value. Subpanel, amount of training (epochs) until 0.8 similarity to final weights is reached for different brain areas. V1, which learns a final representation close to its initialization, is omitted. In our local plasticity model with LW loss (**B**.), even with simultaneous plasticity across the whole brain, connections onto the earlier areas (V1,V2) converge towards their final values earlier than connections to higher areas (temporal cortex). Training time necessary to reach the 0.8 similarity threshold monotonically increases with depth. Contrariwise, with non-local plasticity (**B**., BackProp through the whole brain from the same loss applied to the IT output only), connections to all areas (except V1) evolve in parallel; all areas reach 0.8 similarity almost synchronously.

Interestingly, our local learning rule leads to partially sequential learning, even if we do not impose CPs. In a control run, we used the learning rule of Equation 2) in all brain areas simultaneously, such that synapses across all brain areas adjust in parallel (Figure 2B). However, synapses in the model area V1 (area 1 of the visual stream) reach more rapidly values close to their final states (cosine similarity above 0.8, Figure 2B, inset) than synapses in area IT (area 6 of the visual stream). With end-to-end training using BackProp (parallel plasticity), synapses in all areas approach their final values at roughly the same speed (Figure 2C, inset). Thus, staggered learning is a phenomenon that emerges naturally in local but not end-to-end learning.

## 3 Critical periods reduce the energetic cost of learning

Are there advantages to sequential learning with staggered CPs? We hypothesized that the computational advantage of sequential learning consists in a smaller number of weight updates compared to parallel plasticity, for a similar final performance. Since each up-or down-regulation of a synapse has a metabolic cost, a reduction of the number of times a synapse changes could be an evolutionary advantage.

To test our hypothesis, we trained the network for different durations (number of epochs), with the CPs of subsequent areas opening either all at the same time (parallel) or one after the other (staggered). For both conditions, we evaluated the total number of synaptic weight changes during learning, assuming all plastic synapses update after each stimulation (Methods).

We find that indeed, for the same metabolic cost, the sequential model with staggered CPs achieves better accuracy than the parallel model (Figure 3A). However, the benefits of sequential training are no longer observable in the limit of a large number of synaptic updates, as the graphs of both approaches eventually meet (Figure 3A). In addition, we observe a data-plasticity tradeoff: for a given number of synaptic updates, the sequential model uses more data, and for a given amount of data (epochs), the parallel model achieves better accuracy (Figure 3B).

**Figure 3.**
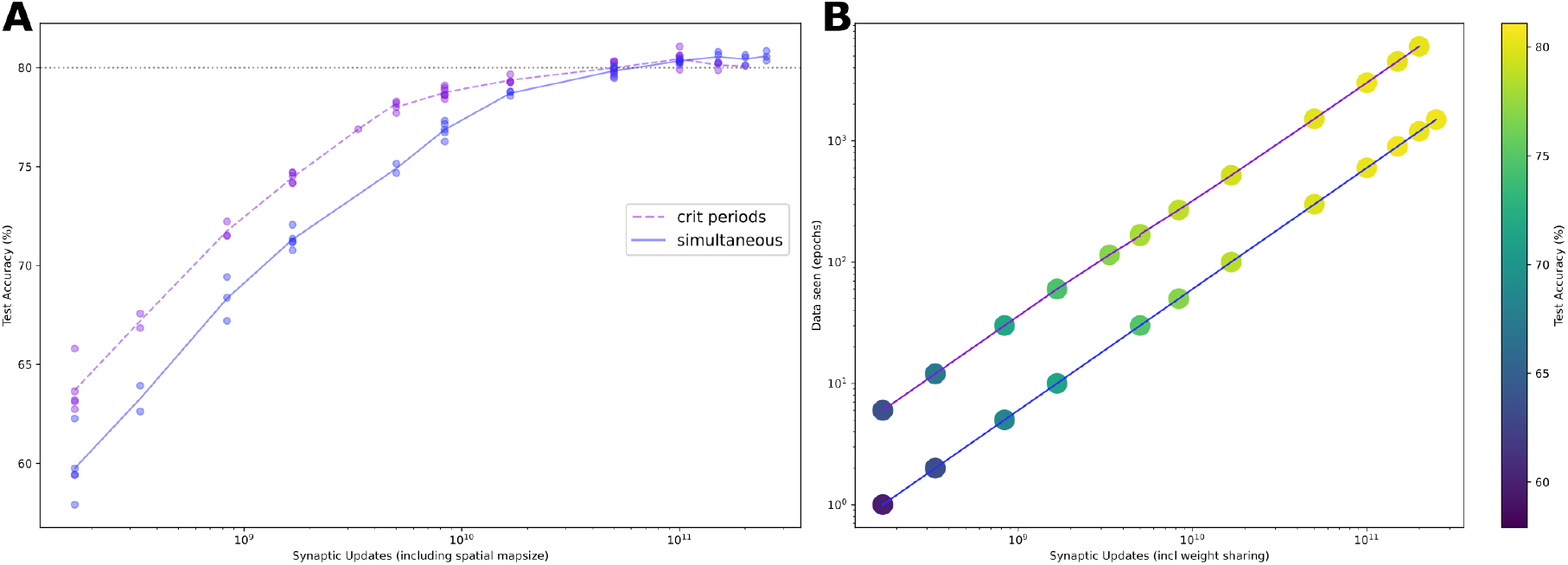
Staggered CP plasticity updates synaptic weights more efficiently than parallel plasticity. **A.** Linear decodability (test accuracy, vertical axis) for parallel (blue) and staggered CP (purple) training as a function of overall number of synaptic updates (log-scale). Lines show average over three to five different random seeds. Sequential training is consistently ahead of parallel training until performance saturates. **B**. Amount of data used for training (y-axis) as a function of synaptic updates. (Test accuracy represented by the color). Here we observe a data/plasticity tradeoff: as seen in panel A, for a given budget of synaptic changes (location along the horizontal axis), staggered CP learning (purple line) reaches better accuracy than parallel learning (blue). But for a given budget of data (location along the vertical axis), parallel learning reaches better accuracy.

## 4 Acquired visual representations are useful for behaviour

While classic supervised learning combined with the Backpropagation Algorithm is designed to optimize the behavior for a given task, self-supervised learning comes with no intrinsic guarantee of optimality for a given task. We investigated whether the visual representation that develops during learning with Eq. (2) might, even though potentially suboptimal for any given task, offer *generic* utility for diverse behavioral tasks (*i*.*e*., this visual representation is a good basis to learn multiple tasks from). To test this hypothesis, we performed several reinforcement-learning (RL) experiments similar to experiments conducted on mice. In all cases, we apply a biologically interpretable RL algorithm [88, 89] based on eligibility traces (Methods), to maintain biological plausibility throughout the entire learning pipeline - from representation to decision making. The representation of the input (i.e. the ‘state’ in the sense of RL) is always the one acquired with our learning algorithm during exposure to thousands of images from the STL-10 data base during a sequence of staggered CPs.

We first tested visual navigation in a T-maze, in the absence of any input from path-integration. The arms of the maze are distinguishable since their walls are densely covered with images from the STL-10 database. The RL agent learns to navigate to the reward located at the end of one of the arms. The only input it receives is the egocentric view of the maze from its current location, encoded by our model in areas 4-6 of the visual stream. As shown in Supplementary Figure 9 of [90], the agents learn to solve the task using the existing visual representations provided by our previously acquired model without the need of further fine-tuning of synapses in the visual stream.

The second artificial task is inspired by the Morris water maze, but is otherwise similar to the first one. The agent is placed in a circular room with concrete walls. The only available landmarks are a few isolated “posters” (seven randomly chosen STL-10 images) on the walls. Similarly to the water maze with a single escape platform in opaque water, there is a single reward location that is not visible. Consistent with experiments [91], artificial agents can learn this task (Figure 4A).

**Figure 4.**
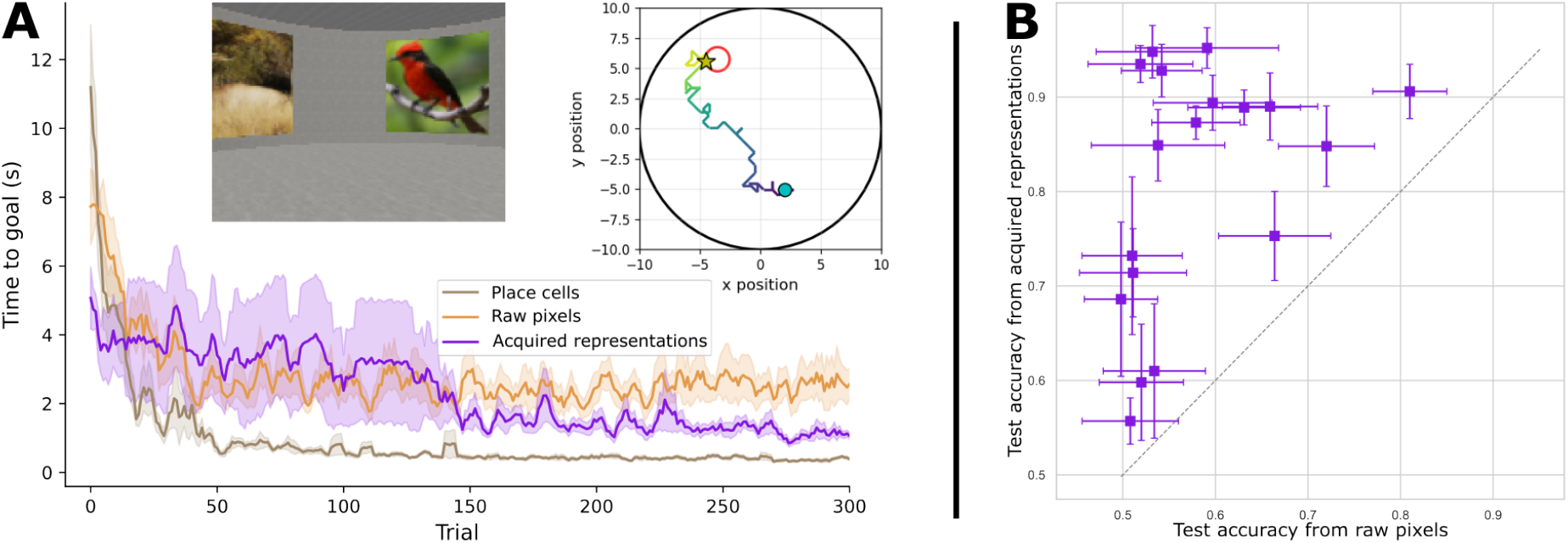
Acquired visual representations are useful for behaviour. **A.** Morris watermaze task: the agent learns to navigate to a hidden target at a constant location. Top left, example egocentric view. Top right, example trajectory in the maze (goal is the red circle). Bottom, learning curves for an RL agent whose state is either the representation by our learnt visual system of its egocentric view (purple), the raw egocentric view for comparison (orange), or the exact position encoding by place cells (brown). **B**. Binary cue-guided bandit task: the agent learns to choose one of two actions depending on the cue category. Success rate of the agent using our learnt visual system (y-axis) *vs* agent receiving the raw image, for different tasks (category pairs). The cues are images from the large dataset ImageNet, and the cue category pairs are chosen to cover varying degrees of difficulty (e.g. easy: banana *vs* unicycle; hard: laptop *vs* computer keyboard). Our learnt visual representations systematically improves the separability of categories (all points above the diagonal), including for hard tasks where the agent behaves randomly if receiving raw pixels input (points on the left, around 0.5 test accuracy for the pixels-based model). The tasks shown are the following: laptop *vs* computer keyboard / laptop *vs* monitor / monitor *vs* television / tiger shark *vs* hammerhead / common iguana *vs* green lizard / green lizard *vs* sea snake / sea snake *vs* hammerhead / sports car *vs* convertible / traffic light *vs* sports car / vending machine *vs* grocery store / tobacco shop *vs* grocery store / trombone *vs* vending machine / dam *vs* vending machine / teddy *vs* vending machine / teddy *vs* dam / cello *vs* teddy / laptop *vs* vending machine / banana *vs* unicycle.

The last RL experiment is a family of visual binary decision making tasks, called bandit tasks in the RL literature. The agent is located in front of a screen and must choose one of two actions depending on a visual cue. The visual cue is an image from the ImageNet dataset. If the image belongs to class A (B), the correct action is a1 (a2). Neither of the two classes is part of the STL-10 labeled set, such that the task requires generalization of the learned representations. Different tasks are defined by different class pairs, and are of varying difficulties. The bio-plausible RL agent learns to discriminate the classes and choose the appropriate actions, with varying degrees of success depending on class separability, but always above chance level. Relatively poor performance for hard class pairs is expected given that our model was trained on the much easier dataset STL-10, and is known to still lag far behind standard BackProp models on harder tasks like ImageNet. However, on many tasks that are hard (task accuracy at chance level for the agent based on raw images rather than learnt representations), the encoding by our acquired visual stream provides great improvement.

Overall, with these experiments, we show that the representations learned by our model are useful for bio-plausible learning of navigation or decision-making behaviours. This finding is consistent across a variety of visual input, relying on within-distribution (STL-10) and out-of-distribution (ImageNet ) images, presented either in frontal view (last experiment) or in perspective, embedded in a spatial view (maze tasks). Thus, our model learns meaningful, actionable and generalizable representations fit to support future behavioral learning in hitherto unknown tasks.

## 5 Discussion

Our findings on the relationship between sequential critical periods plasticity and local learning raise many interesting hypotheses and questions.

### Biological learning

Critical periods *are* found to be staggered across the hierarchy in the cortex of mammals. In light of our findings that

- sequential learning can be more metabolically efficient,
- sequential learning obstruct BackProp learning but empowers local learning,

the existence of sequential critical periods in mammals suggests greater similarity of their biological learning mechanisms with local learning than BackProp learning, consistent with current opinions on biological learning. Our model additionally formulates the hypothesis that sequential learning can constitute an evolutionary advantage in the form of metabolic economy while growing up.

However, our model of sequential learning and the corresponding results hold because of the strictly feedforward architecture of the network. In the more general case of feedback recurrence, as is undeniably found in the brain, the staggering of critical periods looses its edge, as seen with BackProp learning. This raises the interesting question of feedback in biological critical periods, which potentially interacts with the question of remaining plasticity: while critical periods are well-defined for specific features, significant plasticity undeniably persists in the adult brain. Perhaps the feedback connections, in the visual system, are matured later than early-stage processing, and can contribute to fine-tune representations?

### Translational perspectives: re-opening critical periods after lesion

Strokes, a major health problem affecting 20% of the population in the course of their life, induce localized lesions in the brain. It has been suggested that future pharmacological methods in combination with stimulation of the brain area could ‘re-open’ a critical period so as to enhance brain plasticity and contribute to compensation of associated problems [71, 92, 93]. Our model of bio-plausible learning with sequential critical periods opens an avenue for modeling such interventions *in silico*.

### Efficient SSL

Finally, the combined properties of local learning, efficiency and single-layer learning also hint at potential applications in neuromorphic computing, energy-efficient ML, and distributed learning or with small memory available.

## 6 Methods

### 6.1 Implementation details for our model of local learning

Our algorithm relies on the learning rule CLAPP [72]. Here we explain how we use and interpret it; details that are not mentioned here are identical to the original setup of [72].

#### The generalized Hebbian learning rule

Here we provide implementation details complementing the model description in the main text. The non-linearity *ρ* in Eq. 1 is the ReLU function. The convolution kernels are of size 3×3, following measures of the scaling of receptive fields of cells from one area to the next along the visual pathway [94].

In the plasticity rule (Eq. 2), the first post-synaptic term is the derivative of the somatic activity, post_*i*_(*t*) = *ρ*′(∑_*j*_ *w*_*ik*_pre_*k*_ ). The learning rate *η*(*t*) is layer-specific and integrates a global self-supervised signal *y* ∈ {−1, +1} and a layer-specific surprise signal *H* ∈ {0, +1} that vanishes and suppresses plasticity when the layer is already fit. The surprise signal *H* emerges from the derivation of the layer-specific Hinge loss (Eq. 3) [24, 72] and is shared across all neurons in a given layer. Positive and negative samples alternate in equal numbers: an image is presented and plasticity matches the lateral predictions (*y* = +1), then the image switches and plasticity contrasts activity and prediction (*y* = −1).

#### Spatial architecture

As in [72], the input to the CNN is a patch of an image, rather than the whole image. However, the overall design differs in several aspects. First, we use a patch size of 27×27 pixels (instead of 16×16 in [72]), which enables the effective receptive field (RF) size to keep growing after layer 4 (given the convolution architecture, the RF of neurons in the 4th convolutional layer is 14 and that of the 5th layer is 24). We found that this improved the representation quality in higher layers (data not shown). The cropping augmentation of STL-10 images (before cutting out the patches) is set to 92×92 (randomly for training, in the center for testing).

To represent lateral connections across the retinotopy, we apply the CLAPP loss to the representations of a pair of neighbouring patches: *z* is the spatially-averaged neural activity for a patch at a given convolutional layer, and *c* is the same for an adjacent patch. The apical prediction weights *W*^*pred*^ are direction dependent (there are four sets of apical weights per convolutional layer: one for *c* from each cardinal direction). We apply all four losses (or those available, if *z* is from an edge patch) simultaneously.

#### Computation of the number of synaptic weight updates

We take into account that our model is a convolutional neural network by ignoring the weight sharing and calculating the number of synaptic connections as a function of the spatial map size: 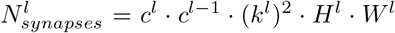 where *c*^*l*^ is the number of channels/features of layer *l, k*^*l*^ = 3 is the kernel size of layer *l* and *H*^*l*^, *W*^*l*^ are the height and width of the feature map at layer *l*. Note that the actual weight vector is only of size *c*^*l*^ · *c*^*l*−1^ · (*k*^*l*^)^2^ due to convolutional weight sharing. In practice, the mapsize factor (which differs from each layer) does not impact the results shown here. In Figure 3A we represent the sequential model with a dashed rather than full line to highlight that points with more synaptic updates *cannot* be reached by continuing the training from points to the left, due to the strict sequentiality. For parallel critical periods, intermediate training durations constitute intermediate points.

### Critical periods

We define the critical period (CP) of a brain area as the epoch(s) during which it is plastic. Plasticity of incoming weights to a brain area is inactivated outside of its critical period. In our model of parallel plasticity, all CPs cover the whole training. In our model of staggered plasticity, CPs open sequentially across the hierarchy, the CP of a brain area only opening after the CP of the upstream area has closed. Note that in the model of BackProp learning with sequential CPs (Figure 2), the activities and gradients flow through all layers regardless of the timing, but during any epoch only the connections of the one plastic area update. In the layerwise (LW) models, only the activities flow (forward) through frozen areas.

Sequential training enables training each area to a different degree. We found that the usefulness (to higher layers) of the representations learned in V1 plateaus after roughly 20 epochs, but no such clear effect was visible for the higher layers. Therefore, in our models of staggered plasticity, the critical periods of all areas last equal numbers of epochs, except the first which lasts no more than 20 epochs.

### 6.2 Evaluating and testing

#### Classification on STL-10

Following standard methods [72, 53], we evaluate the quality of the self-supervised representations by training (for 600 epochs) a linear readout layer to perform classification on the labeled part of STL-10, and report accuracy on the validation set. The colored data points in Figure 1B correspond to a single-area representations: the input to the linear classifier is the concatenation of the (spatially averaged) neural activities *z* in one brain area (convolutional layer) for all patches. The brown crosses correspond to mutliple-area representations, which are obtained by concatenating the single-area representations of three consecutive areas. For the Back-Prop model, considering that representations in all areas are optimized *for the final representation*, we use only the neural activities of the last area.

#### Statistical comparison to other published models

In Figure 1C., we show the p-value for a Welch T-test (`scipy.stats.ttest_ind_from_stats`) based on 6 different random seeds for our model, and the reported average, number of samples, and standard error of the mean in [53] and [79]. The p-values are Holm-Bonferroni corrected for multiple testing. No variability metric is reported in [72].

### 6.3 Reinforcement learning experiments

To test whether the representations learned by our model support biologically relevant behaviour, we designed two reinforcement learning (RL) tasks:spatial navigation in a watermaze, and a visual binary bandit. In both tasks, the RL agent receives as input a visual observation encoded by the frozen representation network (our CLAPP-based model or, as a control, a SimCLR encoder), and learns to act using a lightweight actor-critic algorithm. The encoder weights are never updated during RL training; only the small actor and critic weight vectors are learned.

These tasks complement that in the Supplementary Figure of [90], which already tested the representations of our model with sequential critical periods in a RL setting.

#### Common RL algorithm: Actor-Critic with eligibility traces

All tasks use the same actor-critic learning algorithm. The agent maintains a critic weight vector **w**_*c*_ ∈ ℝ^1×*D*^ and an actor weight matrix 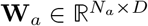, where *D* is the feature dimension and *N*_*a*_ the number of actions.

The value estimate is *V* (*s*) = **w**_*c*_ · **f** (*s*), where **f** (*s*) ∈ ℝ^*D*^ is the feature vector of state *s*. The policy is a softmax over action logits: *π*(*a*|*s*) = softmax(**W**_*a*_ **f** (*s*))_*a*_, from which actions are sampled stochastically.

At each time step, the temporal-difference (TD) error is

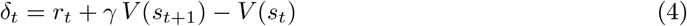

where 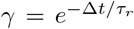 is the discount factor derived from a reward time constant *τ*_*r*_ and simulation time step Δ*t* = 0.01 s. Eligibility traces for the critic and actor decay exponentially:

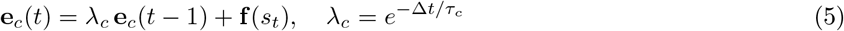

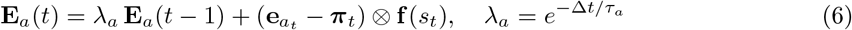

where 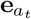 is the one-hot action indicator, ***π***_*t*_ is the softmax policy vector, and ⊗ denotes the outer product. The trace time constants are *τ*_*c*_ = *τ*_*a*_ = 0.5 s. Weights are updated online at every step: **w**_*c*_ ← **w**_*c*_+*α*_*c*_ *δ*_*t*_ **e**_*c*_(*t*) and **W**_*a*_ ← **W**_*a*_+*α*_*a*_ *δ*_*t*_ **E**_*a*_(*t*), where *α*_*c*_ and *α*_*a*_ are the critic and actor learning rates respectively.

This learning algorithm, taken from [95], was chosen because it aptly complements our bio-plausible generalized three-factor representation learning rule with a bio-plausible three-factor reinforcement learning rule.

#### Training and hyperparameter tuning

For each task and for each encoding type, we run a small grid search (in steps of a half order of magnitude, 3 · 10^−*n*^, and with 3 different random seeds for each) on the learning rates of the actor and the critic; here we report findings for the best learning rates found.

We train ≥ 10 agents independently with different seeds for each case and show the aggregated results.

#### Visual encoding

In all tasks, the agent’s visual observation (an RGB image) is encoded by a frozen feature extractor. We compare:

- Our model: the six-layer convolutional network described above, trained on STL-10 with critical periods. The agent’s input is the neural representation obtained from the highest three areas, as described in subsection 6.2.
- raw pixels: no encoder, the agent’s input is directly the visual observation (pixel values in [0 − 1]).

The visual features are *L*^2^-normalized before being passed to the actor-critic.

To avoid repeated forward passes through the encoder during RL training, visual features are precomputed (in the spatial task, for a dense grid of agent positions and viewing directions in the environment), stored on disk, and retrieved at runtime via nearest-neighbor lookup.

#### Task 1: Morris watermaze

##### Environment

The environment is a circular room built in the Miniworld 3D simulation framework. The room is a regular polygon with 100 wall segments approximating a circle of radius *R* = 10 m. Walls have a uniform concrete texture and a height of 5.48 m. A set of *N*_*p*_ STL-10 images (“posters”) are placed on the walls as the sole visual landmarks. The posters are evenly spaced around the perimeter and collectively cover a fraction *c* of the total perimeter (default *N*_*p*_ = 7, *c* = 0.5). The floor uses a concrete texture and the agent’s camera produces RGB observations of 160 × 120 pixels.

At the beginning of each session, a goal position **x**_goal_ is drawn uniformly at random inside the room (within an effective boundary of radius 9.9 m), and remains fixed for the entire session. The goal is invisible; the agent must learn to navigate to it using only visual observations. The goal is considered reached when the agent is within *r*_goal_ = 1.0 m of **x**_goal_.

As in an experimental Morris watermaze experiment, the reward is not visible, and its location is kept constant across training (different across random seeds).

Note that in this environment, the landmarks are from the same dataset as our model was trained on (but the input view is a rendering of the maze in a given direction).

##### Agent state representation

The feature vector **f** (*s*) is the visual encoding of the current observation (1024-dimensional for CLAPP pooled, 2048 for SimCLR). As a baseline, we also test place-cell-only agents using 169 Gaussian place cells on a Cartesian grid (spacing 2.0 m, width 1.0 m, peak rate 20 Hz) with no visual input.

##### Action space and movement

The agent uses a discrete action space with 8 possible actions, corresponding to a step in each of the four cardinal and four diagonal directions.

##### Reward and trial structure

Each trial begins at a uniformly random position inside the room. The agent receives a reward of *R*_goal_ = +100 upon reaching the goal and a per-step penalty of *R*_time_ = −1. The value function is set to zero at the goal (terminal state). Each trial lasts at most *T*_max_ = 20 s (2000 time steps at Δ*t* = 0.01 s).

##### Precomputed features

Visual features are precomputed for maze views on a grid of 50 ×50 positions, with a wide field of view and facing north. At runtime, the nearest grid point to the agent’s current position is found and the corresponding precomputed representation of the corresponding view is used as the state.

##### Hyperparameters

In the watermaze experiment, we apply exponential learning rate decay with a factor of 0.995 (applied between trials).

For our model, hyperparameter search found best reward time constant *τ*_*r*_ = 1 and base learning rates *α*_*c*_ = 1 × 10^−7^ and *α*_*a*_ = 1 × 10^−3^. The agent receiving raw pixels input has *τ*_*r*_ = 0.1 and base learning rates *α*_*c*_ = 1 × 10^−5^ and *α*_*a*_ = 1 × 10^−4^. The agent based on place cells has *τ*_*r*_ = 2 and base learning rates *α*_*c*_ = 1 × 10^−5^ and *α*_*a*_ = 1 × 10^−5^.

#### Task 2: Visual binary bandit

##### Task

This is a one-step decision task. On each trial, the agent is shown a single image belonging to one of two visual categories and must choose one of two actions. Action 0 is correct for category A images and action 1 for category B. The agent receives a reward of +1 for a correct choice and 0 otherwise. There is no temporal structure: each trial consists of a single observation–action–reward step.

##### Visual stimuli

Images are drawn from ImageNet. We tune the hyperparameters on the category pair “banana” and “unicycle”, then train and test other categories as shown in Figure 4B. We chose categories that are not among the 10 classes of the STL-10 labeled dataset; therefore this task tests generalization of the learned representations beyond the reported STL-10 accuracy. Images are resized to 160 × 120 pixels. Up to 300 images per category are used for training (from the ImageNet training split); testing uses images from the ImageNet validation split.

##### Agent

Since there is no temporal structure, the update simplifies to one-step TD learning without eligibility traces: *δ* = *r*−*V* (*s*), and the critic and actor updates use the current feature vector directly (no trace accumulation). The actor weight matrix is of size 2 × *D* and the critic weight vector of size 1 × *D*, with Xavier initialization.

##### Hyperparameters

Chosen learning rates are *α*_*c*_ = 5 × 10^−6^ and *α*_*a*_ = 5 × 10^−5^. Each session runs 2000 training trials followed by 100 test trials (frozen agent weights, validation images). We run 10 sessions (different random seeds) for each category pair.

